# The mutual repression between Pax2 and Snail factors regulates the epithelial/mesenchymal state during intermediate mesoderm differentiation

**DOI:** 10.1101/2021.06.30.450631

**Authors:** Juan M. Fons, Oscar H. Ocaña, M. Angela Nieto

## Abstract

The pronephros is the first renal structure in the embryo, arising after mesenchymal to epithelial transition (MET) of the intermediate mesoderm, where Pax2 induces epithelialisation of the mesenchyme. Here we show that, in the early embryo, Snail1 directly represses *Pax2* transcription maintaining the intermediate mesoderm in an undifferentiated state. Reciprocally, Pax2 directly represses *Snail1* expression to induce MET upon receiving differentiation signals. We also show that BMP7 acts as one such signal by downregulating *Snail1* and upregulating *Pax2* expression. This, together with the Snail1/Pax2 reciprocal repression, establishes a regulatory loop in a defined region along the anteroposterior axis, the bistability domain within the transition zone, where differentiation of the neural tube and the somites are known to occur. Thus, we show that the antagonism between Snail1 and Pax2 determines the epithelial/mesenchymal state during the differentiation of the intermediate mesoderm and propose that the bistability zone extends to the intermediate mesoderm, synchronizing the differentiation of tissues aligned along the mediolateral embryonic axis.

**Summary:** The antagonism between Snail and Pax2 factors in the embryonic differentiation zone, tightly regulates the timing of mesodermal epithelialisation

## INTRODUCTION

During embryonic development, most of the tissues are derived from the sequential activation of epithelial-mesenchymal (EMT) and mesenchymal-epithelial transitions (MET) (Thiery et al., 2009). A prototypic example is the development of the sequential renal structures, pronephros and metanephros. The pronephros is the first epithelial structure of the urogenital system that arises from the intermediate mesoderm after MET. It encompasses the nephric duct and nephric cord running along both sides of the embryo adjacent to the somites. It elongates posteriorly until it reaches the metanephric mesenchyme where the adult kidney will develop (James and Schultheiss, 2003)(Dressler, 2009).

Snail transcription factors are potent EMT inducers in the embryo at gastrulation (Acloque et al., 2011) and neural crest delamination (Nieto et al., 1994), among other embryonic processes. During the development of the renal system in vertebrates, they are expressed in the intermediate mesoderm before pronephros epithelialisation and later in the metanephric mesenchyme prior to the MET that leads to the formation of the renal vesicle (Boutet et al., 2006). In both cases, epithelialisation concurs with the repression of Snail factors, which are maintained silent in the adult, although they are reactivated in pathological conditions like fibrosis, where they induce a partial EMT and instruct the interstitium to promote fibrogenesis and inflammation (Grande et al., 2015)(Lovisa et al., 2015). If the reactivation occurs in cancer cells, they disseminate from the primary tumour initiating the metastatic cascade (Ocaña et al., 2012)(Yang et al., 2020). By contrast, *Pax2* is an essential promotor of MET and required for the specification of the renal epithelial lineage (Dressler et al., 1993)(Torres et al., 1995)(Bouchard et al., 2002). It is expressed in the intermediate mesoderm just prior to MET and continues to be expressed in the epithelial derivatives of the pronephros, mesonephros and metanephros (Dressler et al., 1990)(Dressler and Douglass, 1992)(Bouchard et al., 2000)(Bouchard et al., 2002)(Narlis et al., 2007). Another important molecule that promotes epithelialisation and MET during renal development and fibrosis is BMP7, a member of the TGFβ superfamily (Massagué, 2012) expressed in the pronephros, mesonephros and metanephros (Lyons et al., 1995)(Vukicevic et al., 1996). In agreement with this, *Bmp7* null mutant mice display renal hypoplasia with a reduction of the epithelial compartment (Dudley et al., 1995)(Luo et al., 1995). Furthermore, BMP7 treatment prevents and reverses acute renal fibrosis restructuring the epithelial tubules of the kidney inducing MET by antagonizing TGFβ signalling (Zeisberg et al., 2003)(Sato et al., 2003)(Zavadil et al., 2004)(Zeisberg et al., 2005), a well known EMT inducer and *Snail* transcriptional activator (Peinado et al., 2003).

We have examined the relationship between EMT and MET inducers Snail, Pax2 and Bmp7 in the intermediate mesoderm, as it is an excellent model to study epithelial plasticity in the embryo and found that a reciprocal repression between Snail1 and Pax2 controls the timing of differentiation. This negative regulatory loop is triggered by Bmp7, which simultaneous represses Snail1 and activates Pax2 expression. This loop is activated in the region were differentiation of the neural tube and the paraxial mesoderm also occurs, indicating that it is integrated in the synchronization of differentiation processes along the mediolateral axis.

## RESULTS AND DISCUSSION

### *Snail1* and *Pax2* are expressed in mutually exclusive domains in the developing mesoderm

Given that the silencing of *Snail1* expression correlates with the epithelialisation of the metanephros (Boutet et al., 2006) and that *Pax2* is required for the epithelialisation of pro and metanephros (Bouchard et al., 2002)(Narlis et al., 2007), we decide to compare the expression of both factors at early stages of mesoderm development in the chick embryo (Fig. 1). *Snail1* is expressed in mesenchymal tissues including the undifferentiated intermediate and lateral mesoderm (Fig.1A-C) but absent from the nephric duct and cord of the pronephros (Fig.1B). By contrast, *Pax2* is expressed in the epithelial structures of the pronephros but not in the undifferentiated intermediate mesoderm (Fig.1D-F). We performed a double FISH and confirmed the absence of *Snail1/Pax2* co-expression (Fig.1G-I). These results show that *Snail1* and *Pax2* expression domains are mutually exclusive in the developing embryo and, in particular, in the differentiating intermediate mesoderm.

**Fig. 1.**
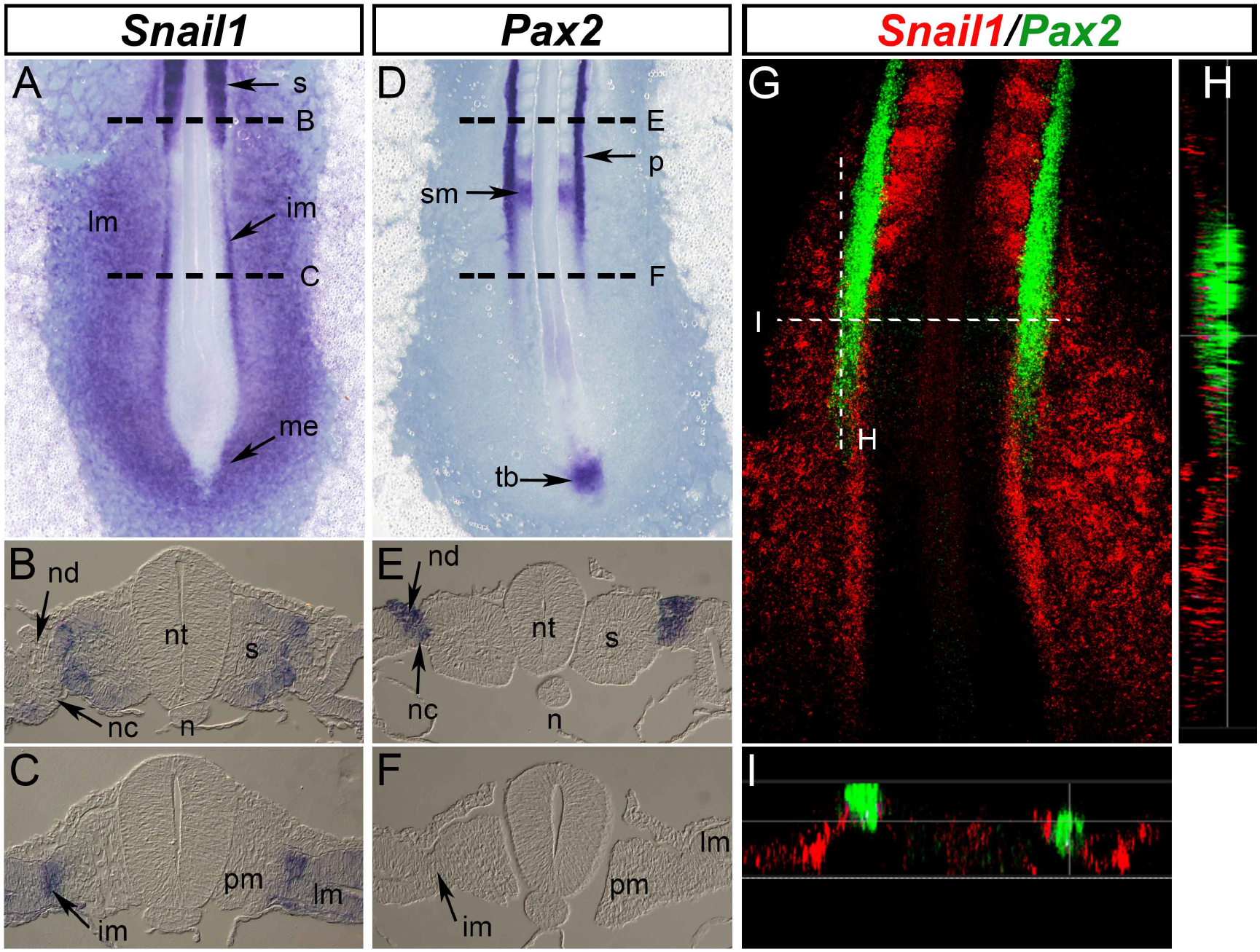
*Snail1* and *Pax2* are expressed in a mutually exclusive pattern in the developing embryo. Dorsal view of chicken embryos at HH11 and their transversal sections. (A-C) *Snail1* is expressed in the ingressing mesendoderm (me), undifferentiated intermediate mesoderm (im), lateral plate mesoderm (lm) and somites (s) (n=10). (D-F) *Pax2* is expressed in the pronephros (p), somitomeres (sm) and in the tail bud (tb) (n=10). (G) 3D reconstruction of a double *in situ* hybridization (FISH) for *Snail1* (red) and *Pax2* (green) (n=3). Longitudinal (H) and Transversal (I) optical sections from the indicated level in a representative embryo in G. Note the mutually exclusive expression pattern. n: notochord, nc: nephric cord, nd: nephric duct, nt: neural tube, pm: paraxial mesoderm.

### Snail1 prevents epithelialisation of the intermediate mesoderm by directly repressing *Pax2* transcription

The expression pattern of *Snail1* and *Pax2* and their described roles in EMT/MET regulation already suggests a possible genetic interaction. To address this, we first performed gain of function experiments by co-electroporating plasmids coding for Snail1 and GFP and analysed *Pax2* expression (Fig. 2A-D). The regions of the intermediate mesoderm with *Snail1* ectopic expression, (Fig. 2A,B green), showed a downregulation of *Pax2* expression, when compared with the control electroporation and the contralateral side (Fig. 2C,D arrow and asterisk; Fig. S1A-D). Analysis of the *Pax2* promoter showed an E box compatible with Snail1 binding (Cano et al., 2000) (Fig. 2E). Luciferase assays in the *Pax2*-positive human renal embryonic epithelial cells HEK293T (Tamimi et al., 2008) show that Snail1 can repress *Pax2* promoter activity (Fig. 2F) in a dose-dependent manner (Fig. S2A-C). Furthermore, the mutation of the E box into a sequence to which Snail1 does not bind (Batlle et al., 2000) restored promoter activity (Fig. 2F). Chromatin immunoprecipitation assays from chick embryo tissues obtained after electroporation of a plasmid encoding a Myc-tagged Snail1 version (Acloque et al., 2011) confirm the enrichment for Snail binding to the *Pax2* promoter (Fig. 2G), indicating that Snail1 behave as a direct repressor of *Pax2* expression *in vivo*. Thus, *Snail1* expression can maintain the intermediate mesoderm in an undifferentiated and mesenchymal state by preventing Pax2 expression and with that, the premature differentiation of the pronephros. This is in agreement with the finding that the intermediate mesoderm fails to become epithelial and remains mesenchymal in Pax2 mutant mice (Torres et al., 1995). As *Snail1* endogenous silencing in the metanephric mesenchyme correlates with MET and renal differentiation (Boutet et al, 2006), and differentiation relies on *Pax2* upregulation (Narlis et al., 2007), it is likely that *Snail1* also prevents premature metanephros differentiation by repressing *Pax2* expression, in addition to its previously described role in repressing *HNF1β* and *Cadherin 16* expression (Boutet et al., 2006).

**Fig. 2.**
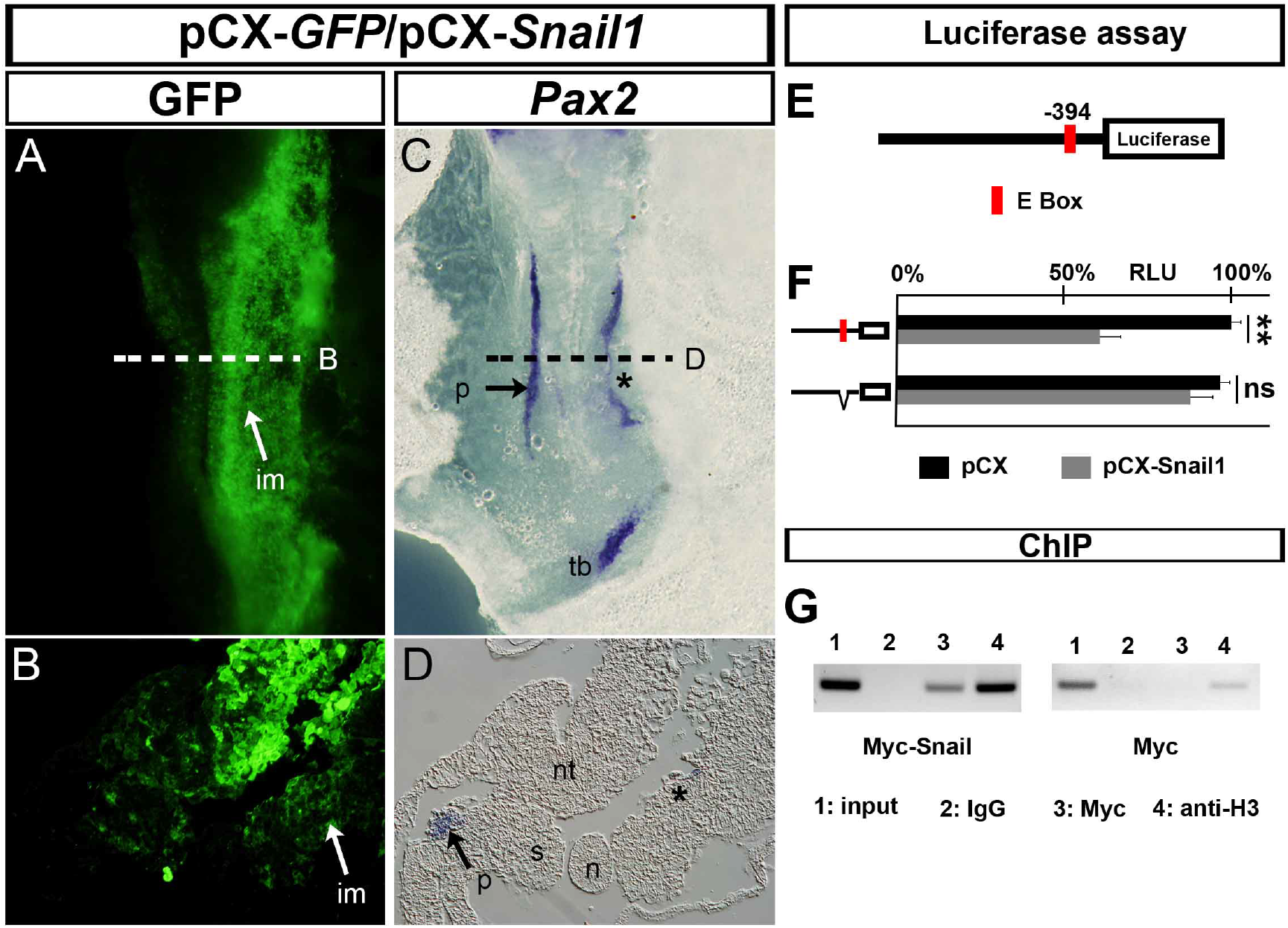
Snail1 is a direct repressor of *Pax2* transcription. (A-D) Chick embryos were co-electroporated on the right side with a vector encoding GFP (pCX-*GFP)* and another containing the *Snail1* coding region (pCX-*Snail1*). Dorsal views and their respective sections showing GFP immunofluorescence (A,B) and *Pax2* expression (C,D). *Snail1* ectopic expression (GFP in A,B) represses endogenous *Pax2* expression (C,D) in the intermediate mesoderm (A, B, white arrows; C, D, black asterisks) (n=9/9). (E) Scheme of *Pax2* promoter fragment (1.8 kb upstream of TSS) cloned into pGL3-Luc vector. A Snail1 binding site (E box) is showed in red. (F) Luciferase assay in HEK293T cells transfected with a vector containing a fragment of the *Pax2* promoter bearing the wildtype or a mutated version of the E box plus either pCX (black bars) or pCX-*Snail1* (grey bars) vectors. Snail1 decreases *Pax2* promoter activity and the mutation of the E box significantly restores the activity (n=3, average representation. T-Test; pValue<0.01; two tails, unequal variance). (G) ChIP assay in chick embryos electroporated with a Myc-tagged Snail1 coding plasmid (Myc-Snail1) or an empty Myc plasmid (Myc) (n=3, representative experiment shown). (1) input; (2) IgG antibody (IgG), negative control; (3) Myc antibody (anti-Myc); (4) Histone3 antibody (anti-H3), positive control. There is amplification of DNA fragments containing the E box in the presence of Myc-Snail1. Therefore, Snail1 represses *Pax2* transcription through direct binding to its promoter. im: intermediate mesoderm, n: notochord, nt: neural tube, p: pronephros, s: somites. tb: tail bud. ** pValue <0.001.

### A Snail1/Pax2 reciprocal inhibitory loop regulates the differentiation of the intermediate mesoderm

As Pax2 is required for MET and renal differentiation (Rothenpieler and Dressler, 1993)(Torres et al., 1995)(Bouchard et al., 2002) and *Snail1* silencing is essential for pronephros differentiation (Fig. 2), we wondered whether Pax2 promoted MET in the intermediate mesoderm at least in part by repressing *Snail1* transcription. We performed gain of function experiments for Pax2 as described above for Snail1 (Fig. 3A-D; Fig. S1E-H). The areas of *Pax2* ectopic expression (Fig. 3A, green) are devoid of *Snail1* transcripts (Fig. 3A-D, asterisks indicate the effect on the intermediate mesoderm). Interestingly, the electroporated mesoderm shows cellular aggregates compatible with a MET process (Fig. 3A, arrowheads), which was confirmed by the re-expression of E-cadherin in the electroporated area (Fig. S3). These results show that the sole activation of Pax2 expression in Snail1-positive intermediate and lateral mesoderm is sufficient to induce epithelialisation. Thus, we examined whether Pax2 could also act as a *Snail1* repressor. We identified a Pax2 binding site (GGGCATGG) 116bp upstream of the Snail1 TSS (JASPAR database) and a luciferase assay in a primary culture of chick embryonic fibroblasts showed that a fragment of 150 bp upstream the TSS co-transfected with a Pax2-encoding vector was sufficient to repress *Snail1* promoter activity in a dose-dependent manner (Fig. 3 E,F and Fig. S2D-F). Deletion of the Pax2 binding site impaired this repression (Fig. 3F). We next performed ChIP analysis in chick embryo tissues and confirmed that Pax2 can bind to the *Snail1* promoter *in vivo* (Fig. 3G).

**Fig. 3.**
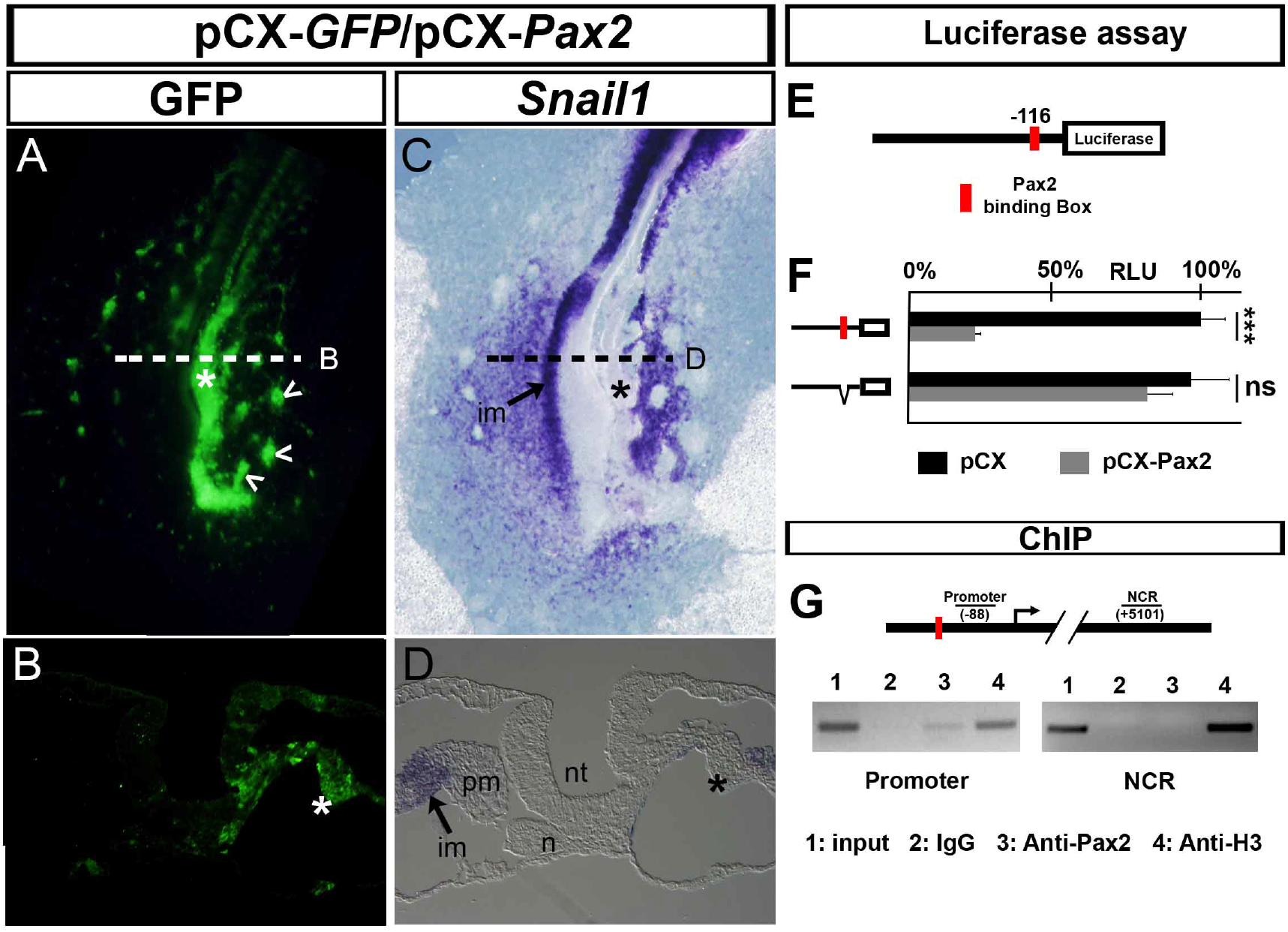
Pax2 is a direct repressor of *Snail1* transcription. (A-D) Electroporation on the right side of a chick embryo with pCX-*GFP* plus pCX-*Pax2* vectors showing IF for GFP (A,B) and ISH for *Snail1* (C,D). (A,C) Dorsal views. (B,D) Sections taken at the level of the dashed lines. *Pax2* ectopic expression (GFP) represses *Snail1* expression in the intermediate mesoderm (asterisks) (n=18/18). (E) Scheme of *Snail1* promoter fragment (0.15kb upstream of TSS) cloned into pGL3-Luc vector. A Pax2 binding box is shown in red. F) Luciferase assay in CEF transfected with a *Snail1* promoter vector containing or not the Pax2 binding box plus pCX (black bars) or pCX-*Pax2* (grey bars) plasmids. Pax2 decreases *Snail1* promoter activity and the deletion of the box restores it (n=3, average representation. T-Test; two tails, unequal variance). G) ChIP assay. Scheme showing the PCR fragments analyzed, one in the promoter region and another one from a Non Coding Region (NCR). (1) input; (2) IgG antibody (IgG), negative control; (3) Pax2 antibody (anti-Pax2); (4) Histone3 antibody (anti-H3), positive control. (n=3, representative experiment shown). Pax2 represses *Snail1* transcription through direct binding to the promoter. im: intermediate mesoderm, n: notochord, nt: neural tube, pm: paraxial mesoderm. ***pValue<0.0001

Altogether, these data show that, similar to Snail1 repressing *Pax2* transcription, Pax2 represses *Snail1* transcription by direct binding to its promoter. Thus, Snail1 and Pax2 establish a reciprocal negative loop, a mechanism widely used during development when fate decisions involve binary choices (Zhou and Huang, 2011). This is observed in the subdivision of embryonic territories, including the decision to become trophectoderm or inner cells mass (Cdx2/Oct4) or ectodermal versus mesendodermal (SoxB1/Snail) (Niwa et al., 2005)(Acloque et al., 2011)(Acloque et al., 2012). This reciprocal repression between Snail1 and Pax2 is also likely to occur during metanephros development and may also have a correlate in pathological conditions. *Pax2* reactivation occurs in fibrotic or injured kidneys (Imgrund et al., 1999)(Hou et al., 2018), and this is thought to be an attempt to recover the normal renal epithelial architecture either through MET of the renal mesenchyme or by increasing proliferation of the remaining epithelial cells (Lindoso et al., 2009). We propose that this role of *Pax2* reactivation maybe achieved by counteracting the activation of *Snail1* in renal epithelial cells upon injury (Grande et al., 2015).

### Bmp7 controls pronephros differentiation by repressing *Snail1* and inducing *Pax2* expression in the Bistability Domain

Then we investigated signalling pathways that could be regulating these genes. We focused on BMP members as they induce pronephros differentiation. Specifically, we tested Bmp7, as it is also required for MET and differentiation of the metanephros (Dudley et al., 1995)(Luo et al., 1995)(Vukicevic et al., 1996). *Bmp7* is expressed in the ectoderm, dorsal neural tube, the notochord and in the pronephros (Fig. 4A,B) and we decided to challenge the embryo by administering BMP7 and assessing its impact on *Snail1* and *Pax2* expression.

**Fig. 4.**
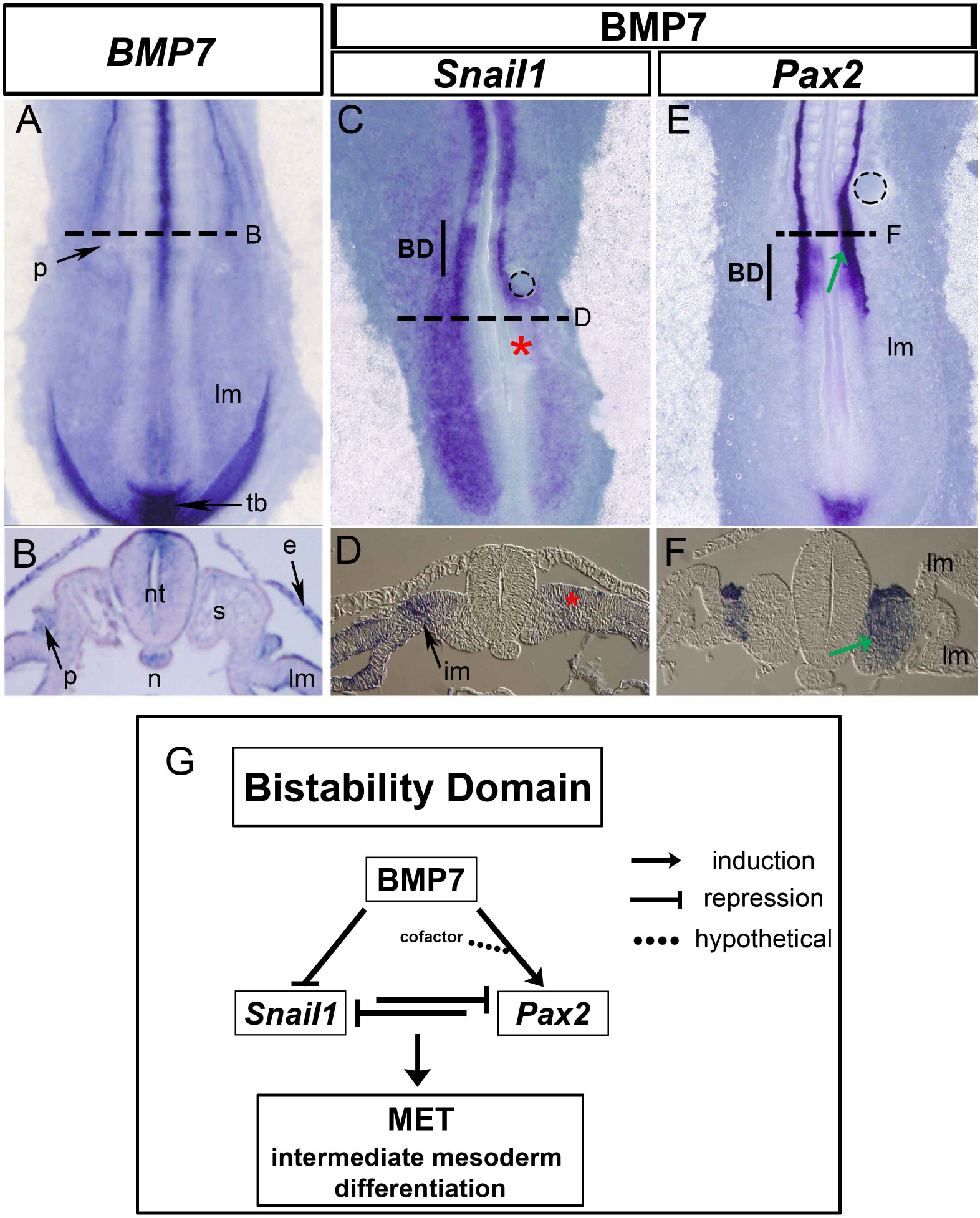
BMP7 represses *Snail1* and induces *Pax2* expression. (A,B) ISH for *Bmp7* and the respective transverse sections. Bmp7 is expressed in the ectoderm (e), dorsal neural tube (nt), notochord (n), pronephros (p) and tail bud (tb). (C-F) Chick embryos bearing a bead (dotted circle) previously soaked in BMP7 (50μg/ml) and subjected to ISH for *Snail1* (C,D) or *Pax2* (E,F). BMP7 represses *Snail1* in the mesoderm (C,D, asterisks,n=3/5) and induces *Pax2* expression in the bistability domain (BD) (E,F, arrows, n=3/5). G) Model for pronephros differentiation. Bmp7 directs the flow of information by repressing *Snail1* and inducing *Pax2* to promote renal differentiation through MET in the intermediate mesoderm within the BD. s: somites, lm: lateral mesoderm, im: intermediate mesoderm.

BMP7 represses *Snail1* transcription in the mesoderm, including the intermediate mesoderm (Fig. 4C,D, asterisks, Fig. S4A-H). In addition, it induces *Pax2* transcription in the paraxial/somatic mesoderm (Fig. 4E,F, arrows, see also Suppl. Fig. 4I-P), while represses Paraxis (Fig. S5). This is consistent with the gradient of BMP signalling in patterning the mesoderm along the mediolateral axis in the chick embryo among other vertebrates (James and Schultheiss, 2005) and with recent data showing that the embryo interprets Bmp7 concentrations thresholds (Greenfeld et al., 2021). Thus, Bmp7 induces pronephros differentiation through two independent mechanisms, by repressing an EMT inducer, *Snail1*, and by inducing a MET inducer, *Pax2* in the intermediate mesoderm. The ectoderm is very likely the source of Bmp7, as ectodermal removal impairs pronephros differentiation (Obara-Ishihara et al., 1999). And again, these data and expression studies suggest that Bmp7 may also control metanephros epithelialisation and differentiation through the same mechanism. In addition, the mechanism behind the attenuation of renal fibrosis by BMP7 (Zeisberg et al., 2003)(Zeisberg et al., 2005) is likely mediated by the repression of *Snail*1 and the induction of *Pax2*.

The ability of BMP7 to activate Pax2 expression in the paraxial mesoderm occurs in a specific region of the anteroposterior axis of the embryo. In addition, we did not observe *Pax2* induction in territories that are naturally incompetent to express it, like the lateral mesoderm, or outside the bistability domain (Fig. 4E,F). This is compatible with the described competence of the cells to respond to BMP according to the embryonic anteroposterior level, synchronizing the differentiation in the different axes (Tucker et al., 2008). As such, the response is within the transition zone (TZ), and restricted to the bistability domain (BD)(Fig. 4E,F, arrows) (Fig. S4I-P). The bistability domain is defined as a convergent area of two opposing gradients within the anterior one-third of the TZ, an anteroposterior gradient of retinoic acid and a posteroanterior gradient of FGF/Wnt (Del Corral and Storey, 2004)(Olivera-Martinez and Storey, 2007)(Goldbeter et al., 2007)(Aulehla and Pourquié, 2010), where neurogenesis (Del Corral and Storey, 2004) and somitogenesis (Aulehla and Pourquié, 2010) occur. Differentiation at the bistability zone is perfectly compatible with the region where binary choices occur (Goldbeter et al., 2007), and particularly with the establishment of reciprocal negative loops as we describe here for Snail1 and Pax2 (Zhou and Huang, 2011).

Interestingly, retinoic acid induces pronephros differentiation (Cartry et al., 2006)(Wingert et al., 2007) and it is known to repress Snail1 during somite epithelialisation (Morales et al, 2007). In addition, Snail1 is downstream of the Wnt/FGF signaling pathway during somitogenesis, and it needs to be repressed for epithelialisation to occur (Dale et al., 2006). Our results show that the bistability domain not only regulates neurogenesis and somitogenesis but also pronephros development. We propose a model for pronephros differentiation (Fig. 4G) in which Bmp7, represses *Snail1* expression and, in combination with other signals, induces *Pax2* transcription in the intermediate mesoderm. *Snail1* maintains an undifferentiated state of the intermediate mesoderm in the posterior part of the embryo, and at the bistability domain, Snail1 and Pax2 stablish a negative reciprocal loop that leads to epithelialisation. Thus, we extend the bistability domain to the intermediate mesoderm, acting as the region in the anteroposterior axis for the initiation of differentiation, synchronizing the different cell populations along the mediolateral axis of the embryo.

## Material and Methods

### Chick embryonic fibroblast isolation

Muscle fragments from HH35 chick embryos were dissected in PBS at 37°C and disaggregated with trypsin for 5 minutes at RT. The homogenize were centrifuged at 1100 rpm for 6 minutes and the pellet were suspended in F-12/HAM and seeded on p10.

### Cell culture

HEK293T cells were cultured in DMEM (Sigma, D6429) with 10% FCS (invitrogen, 10106-169), 1% penicillin/streptomycin (Sigma, P4333) and 1% fungizone (Sigma, A2942). Chick embryonic fibroblasts (CEF) were cultured in F-12/HAM (Sigma, #N6658) with 10% FCS, 1% penicillin/streptomycin plus funginoze.

### Cell transient transfection and luciferase assays

HEK293T cells and CEF were cultured at 10^5^ cells/well in a 12 multiwell plate for 24hr prior to transfection. For luciferase assays, a reaction mix in a final volume of 50μl (for HEK293T) or 100μl (for CEF) in Opti-MEM (Invitrogen, #51985) was made respectively containing 400ng of pGL3basic-*Pax2* promoter or 800ng of pGL3b-*Snail1* promoter vectors and 40ng of CMV-*Renilla* plasmid as endogenous control. To this mix, 50-100-200ng of pCX-*Snail1* or 20-50-100ng of pCX-*Pax2* vectors were added to HEK293T cells or CEF, respectively. Fugene (Fugene, #E2312) was used for HEK293T cells and Lipofectamine (Lipofectamine, #18324-012) for CEF. After adjusting the culture medium to 500μl, the reaction mix for HEK293T was added and cells were assayed for luciferase activity after 48hr. For CEF, the reaction mix was added to 200μl of opti-MEM for 5hr and the medium was replaced with DMEM afterwards. After the transfection, cells were washed twice in PBS and assayed following the manufacturer’s instructions (Promega, Dual-Luciferase Reporter Assay System Part#TM040).

### Embryo electroporation

The electroporation conditions were 5 pulses of 4 volts, 50ms and 1Hz frequency. Embryos at HH3-4 were electroporated with 2 μg/μl of pCX-*Pax2* or pCX-*Snail1* and 0.5μg/μl of pCX-*EGFP* and cultured until HH11 using the easy culture system (Ocaña et al., 2017).

### In situ hybridization

We used a protocol previously described (Nieto et al., 1996) for non-radioactive in situ hybridization. For double fluorescence ISH we followed the method described in Acloque et al., 2008.

### Cytokine treatment

Acrylic beads (Sigma, heparin acrylic beads #H5263) were embedded in a Bmp7 solution (R&D systems, Recombinant Human BMP7 #354-BP) in PBS. Beads were added to embryos at stage HH9 for 8hr before fixing the embryos.

### Gene cloning and mutagenesis

We used 3 chick embryos at HH11 for DNA genomic extraction with chloroform. A 2kb DNA fragment upstream of TSS of *Snail1* promoter (ENSGAL00000008018) was cloned into pGL3 basic plasmid (Promega) using PWO polymerase (Roche, #11644947001) and NheI and HindIII (Biolabs) target sequences flanking the forward (5’-AAAAAAGCTAGCACCGGGTCTACTTGAATTTTG-3’) and reverse (5’-AAAAAAAAGCTTCGTACTCGCCCAGCGCCACCG-3’) primers, respectively. To generate subsequent deletions in the *Snail1* promoter, we used the construct mentioned above as a template and the same restriction enzymes target sequences flanking the primers. The primers used were: 1.2kb fragment forward primer (5’-AAAAAAGCTAGCGAATTACGGCAATTG-3’), 0.6kb fragment forward primer (5’-AAAAAAGCTAGCCCCGCTTCAGTGGGG-3’) and 0.15kb fragment forward primer (5’-AAAAAAGCTAGCTAGTCTGCCCGCCCCGG-3’) with the above described reverse primer.

For the deletion of the Pax2 binding site located at positions -116/-109 from the TSS in the *Snail1* promoter, we used self-complementary primers (forward 5’-CGTCCCATTGGCTCCGGGGGCGGCCC TGCACCGCCCTC-3’, reverse 5’-GAGGGCGGTGCAGGGCCGCCCCCGG AGCCAATGGGACG-3’) without the Pax2 target sequence (5’-GGGCATGG-3’) and carried out whole plasmid amplification. The PCR product was digested with DnpI (Biolabs) for 5hr and transfected into *E. coli* DH5α strain. To subclone the *Pax2* promoter (NM_204793.1) we used a BAC (CH261-43k16) containing the *Pax2* locus. Since the chick *Pax2* promoter sequence was not annotated, we identified a conserved region at *Pax2* locus of *Gallus gallus* by ECR-Browser (http://ecrbrowser.dcode.org) (galGal3 chrUn_random:58143187-58143360), compared this sequence with the chick whole genome by Blast, (http://blast.ncbi.nlm.nih.gov/Blast.cgi Traces- WGS) and we found a contig with a fraction of the chick *Pax2* promoter (Nw_001478025.1). PvuII (Biolabs) and NcoI digestion respectively inside and at the TSS. Allowed us to get a Bac fragment containing the *Pax2* promoter. This was confirmed by southern-blot performed with two α-P32-dCTP labelled probes, one at Nw_001478025.1 (forward primer 5’-AAAGAGACGGAGAAGTGTATTTCG-3’, reverse primer 5’-ACGTTTGAGAAAAACAAAGGAACT-3’) and another one at 5’UTR (forward primer 5’-ATTGCTTTGCTTTGGTTTGTTATT-3’, reverse primer 5’-GAGGCAAAGGAAAGGGAAGA-3’). After isolating the identified band, it was ligated into pGL3 basic plasmid, and transfected into *E. coli* DH5α that were hybridized with the same probe for confirmation, obtaining a 1,7kb DNA fragment upstream of the TSS of the chick *Pax2* promoter. The cloned sequence is the following:

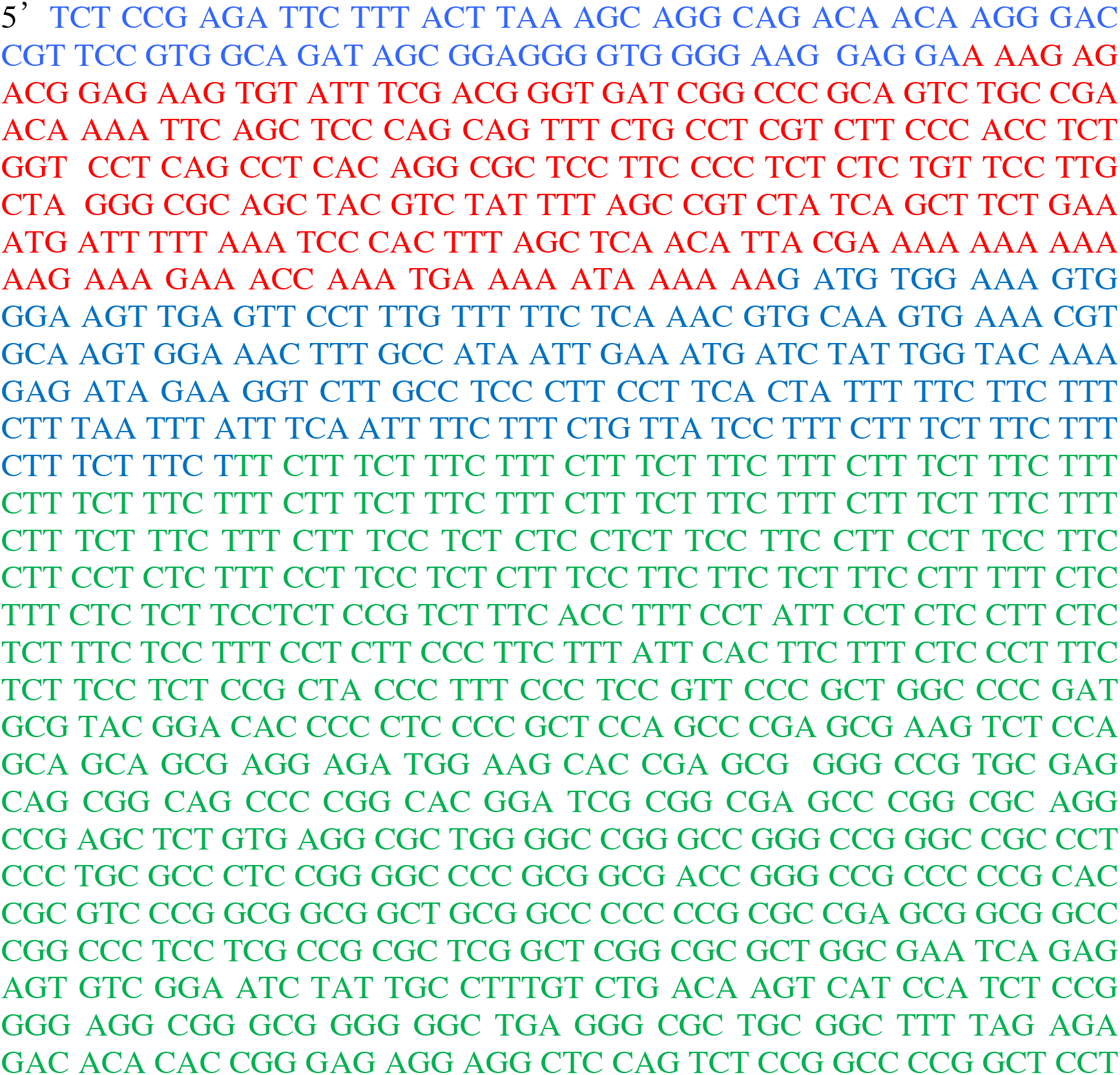

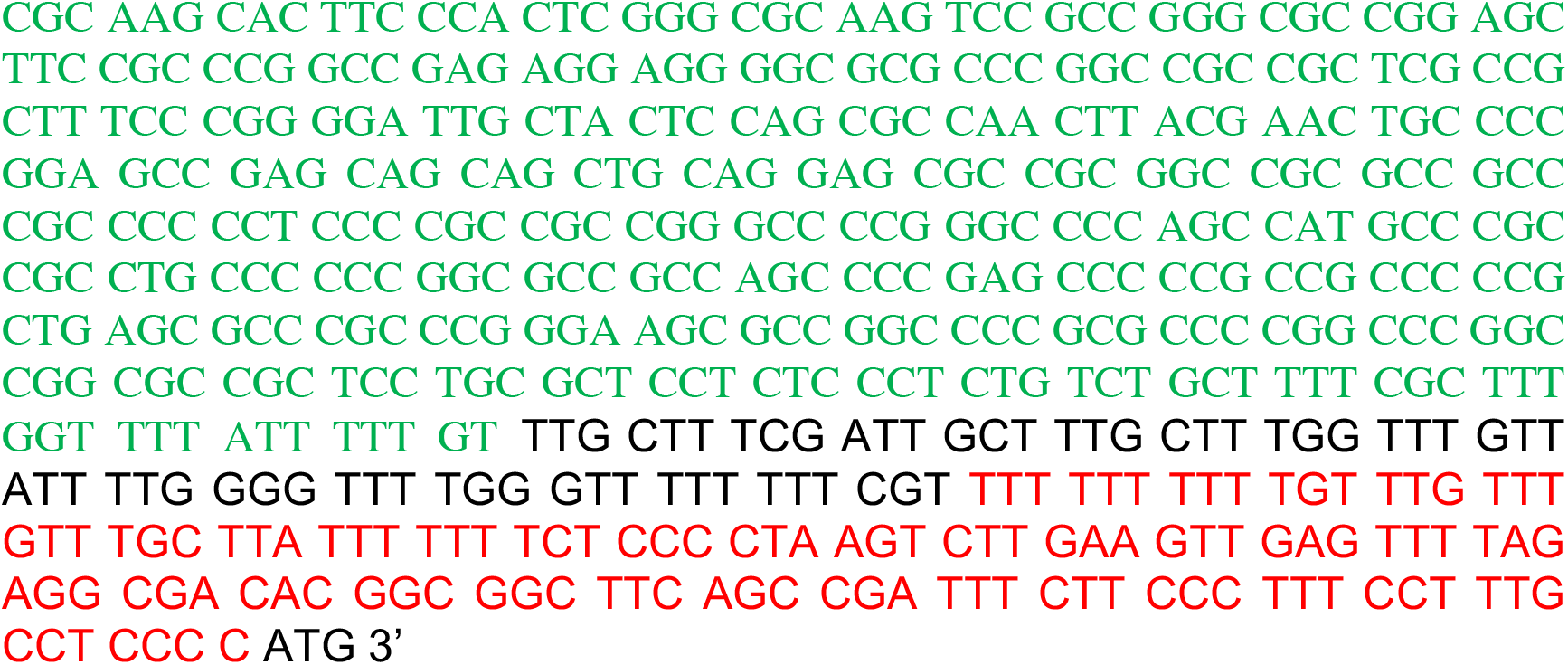

Where black is UTR, Red is probe sequence target, green is new sequence and blue is Nw_001478025.1 contig. Now the sequence of Pax2 promoter has been annotated in ensemble.

For the mutagenesis of the Snail1 binding site (E-box) in the *Pax2* promoter, we mutated the E-box sequence (5’-GCAGCTG-3’ converted to 5’-TTAGCTA-3’) by PCR with the following primers. The primers used were: forward 5’-GCCCCGGAGCCGAGCATTAGCTACAG GAGCGCCGCGGCCG-3’ and reverse 5’-CGGCCGCGGCGCTCCTGTAGCTAATG CTCGGCTCCGGGGC-3’).

### Southern blot

Bacteria containing the BAC (CH261-43k16) were cultured in 200ml of 2YT medium for 16hr and DNA was extracted with a Genomed Biotech Kit (Jetstar plasmid purification maxi kit 220020). DNA was digested with PvuII and NcoI for 6hr. DNA transfer was carried out into a nylon membrane (Milipore, immobilon-P membrane PVDF, IPVH00010) and hybridization was performed with two different probes, one to detect the region upstream of the TSS of *Pax2* promoter (Nw_001478025.1) and another one in the 5’UTR of *Pax2* mRNA. The fragment was amplified with specific primers (see cloning and mutagenesis for the sequence), the PCR product was purified and labelled with α-P^32^-dCTP (Perkin, NEG5134 250uC) following the manufacturer instructions (GE healthcare, radioactive nucleic acid labeling and detection, RPN1633). Hybridization was performed at 65°C o/n and the membrane washed and air dried for 3 minutes before exposure to identify the corresponding band double positive for both probes.

### Chromatin immunoprecipitation

15 embryos at HH11 were treated following the manufacturer’s recommendations (Milipore, Magna ChIP A#17-610) with formaldehyde (Sigma, formaldehyde solution 37% #F1635) and Glycine (Merck #5.00190.1000) with 0.5ml of citoplasmic buffer lysis and 0.3ml of nuclei buffer lysis. The antibodies used were IgG as negative control (2μl, Diagenode #kch-819-015), anti-Histone3 as positive control (2μg, Ab1791), anti-Pax2 (4μg, Ab23799) and anti-Myc (2μg, Ab9132). For Pax2 interaction with *Snail1* promoter, embryos were used without any treatment. For Snail1 interaction with *Pax2* promoter, embryos were electroporated prior to ChIP with 0.5μg/μl of a Myc tagged Snail1 plasmid. The primers used for the amplification of *Snail1* promoter were: forward 5’-CTCCTCGCCCCCCTGTA-3’, reverse 5’-GTACTCGCCCAGCGCCACC-3’, and for the negative control NCR, forward 5’-GCAGCAGCGGCATTATCC-3’, reverse 5’-GTCATGAACCCTTTGGCTTTACC-3’. For the amplification of *Pax2* promoter, forward 5’-GCTACTCCAGCGCCAACTTA-3’, reverse 5’-GGCCGCGGCGCTCCTGCA-3’.

### Immunohistochemistry

Embryos were washed in PBS-Triton X100 at 0.1% and dehydrated in 25%, 50%, 75% and twice in 100% ethanol for 5 minutes at RT. Then, twice in buthanol for 15 minutes and 6 times in wax for 30 minutes at 65°C. Sections were cut at 8 μm and hydrated. A permeabilization step was performed with PBS-Triton X100 at 0.25% twice for 20 minutes. A blocking solution containing PBS-Triton X100 at 0.1% and 10% serum was added for 1hr at RT. Anti-GFP (Santa Cruz #SC9996) antibody was added to the blocking solution at 1/1000 o/n at 4°C. After washing 4 times for 20 minutes at RT with PBS-Triton X100 at 0,1%, a secondary antibody (anti-rabbit-A488, Mol. Probes A-11008) was added at 1/1000 and incubated for 2hr at RT. After washing 4 times for 20 minutes at RT, the slides were mounted with Mowiol (Calbiochem, Mowiol® 4-88 #475904) and imaged.

## Author contribution

J.M.F. performed most of the experiments, analysed and interpreted the data. O.H.O. performed some experiments and with M.A.N conceived the project and interpreted the data. All three wrote the manuscript.

## Acknowledgements

We thank Joan Galceran and Jose Manuel Mingot for helpful discussions and support with the cloning. Sonia Vega for her help and support in managing cell lines, Diana Abad for technical support, Giovanna Expósito for the support at the imaging facility. We also thank Isabel Aller for her help with the southern blot.

